# Full-genome evolutionary histories of selfing, splitting and selection in *Caenorhabditis*

**DOI:** 10.1101/011502

**Authors:** Cristel G. Thomas, Wei Wang, Richard Jovelin, Rajarshi Ghosh, Tatiana Lomasko, Quang Trinh, Leonid Kruglyak, Lincoln D. Stein, Asher D. Cutter

**Affiliations:** Department of Ecology & Evolutionary Biology, University of Toronto; Lewis-Sigler Institute for Integrative Genomics, Princeton University; Department of Pediatrics-Oncology, Baylor College of Medicine; Informatics and Bio-Computing, Ontario Institute for Cancer Research; Departments of Human Genetics and Biological Chemistry, David Geffen School of Medicine, UCLA; Howard Hughes Medical Institute, UCLA; Department of Molecular Genetics, University of Toronto; Bioinformatics and Genomics, Cold Spring Harbor Laboratory; Center for the Analysis of Genome Evolution and Function, University of Toronto

**Keywords:** Population genomics, Caenorhabditis, linked selection

## Abstract

The *nematode Caenorhabditis briggsae* is a model for comparative developmental evolution with *C. elegans*. Worldwide collections of *C. briggsae* have implicated an intriguing history of divergence among genetic groups separated by latitude, or by restricted geography, that is being exploited to dissect the genetic basis to adaptive evolution and reproductive incompatibility. And yet, the genomic scope and timing of population divergence is unclear. We performed high-coverage whole-genome sequencing of 37 wild isolates of the nematode *C. briggsae* and applied a pairwise sequentially Markovian coalescent (PSMC) model to 703 combinations of genomic haplotypes to draw inferences about population history, the genomic scope of natural selection, and to compare with 40 wild isolates of *C. elegans*. We estimate that a diaspora of at least 6 distinct *C. briggsae* lineages separated from one another approximately 200 thousand generations ago, including the ‘Temperate’ and ‘Tropical’ phylogeographic groups that dominate most samples from around the world. Moreover, an ancient population split in its history 2 million generations ago, coupled with only rare gene flow among lineage groups, validates this system as a model for incipient speciation. Low versus high recombination regions of the genome give distinct signatures of population size change through time, indicative of widespread effects of selection on highly linked portions of the genome owing to extreme inbreeding by self-fertilization. Analysis of functional mutations indicates that genomic context, owing to selection that acts on long linkage blocks, is a more important driver of population variation than are the functional attributes of the individually encoded genes.

## Introduction

The record of natural selection in shaping the genetic basis to organismal form and function of a species is inscribed in the genomes of its constituent individuals. Comparisons of genome sequences for each of the related nematodes *Caenorhabditis elegans* and *C. briggsae* have revealed powerful insights into the evolution of functional novelty and constraint in phenotypes and genetic pathways (Cutter et al. 2009; Marri and Gupta 2009; Thomas et al. 2012; Haag and Liu 2013; Verster et al. 2014). The high quality *C. briggsae* genome assembly facilitated such analysis (Stein et al. 2003; Hillier et al. 2007; Ross et al. 2011), and genomic analysis of populations of individuals provides a powerful means to further characterize evolution on contemporary timescales to understand the origins of novelty and constraint (Langley et al. 2012; The 1000 Genomes Project Consortium 2012; Brandvain et al. 2014). Indeed, key questions remain to be solved: how do genomes respond to the simultaneous pressures of mutation, natural selection, and genetic linkage – especially when a novel reproductive mode, facultative self-fertilization, has evolved in the ancestry of a species?

*C. briggsae* is similar to *C. elegans* in many ways, most notably in their streamlined morphology, amenability to genetic and experimental manipulation, and in both being comprised primarily of self-fertilizing hermaphrodites that are found around the globe. But *C. briggsae* is distinctive in having more molecular and phenotypic wild diversity, which is divided along latitudinal phylogeographic lines (Cutter 2006; Raboin et al. 2010; Felix et al. 2013), and by being partly interfertile with its male-female (dioecious) sister species *C. nigoni* (Woodruff et al. 2010; Kozlowska et al. 2012; Felix et al. 2014). Some strain combinations within *C. briggsae* also appear to show reproductive incompatibilities and outbreeding depression (Dolgin et al. 2008; Ross et al. 2011; Baird and Stonesifer 2012). But the extent of genetic exchange and admixture across the genome within this species, as well as a full depiction of its evolutionary history, has remained elusive. Moreover, the extensive linkage disequilibrium conferred on the genome by self-fertilizing reproduction is thought to interact with selection to shape chromosome-scale patterns of genetic diversity (Cutter and Choi 2010; Cutter and Payseur 2013). Consequently, selection, self-fertilization and gene flow all likely interact to control diversity and divergence in ways requiring genomic-scale population information to discern.

These features make *C. briggsae* a powerful tool for dissecting evolutionary pattern and process in connection with trait divergence in nature, especially in combination with its deep experimental toolkit (Koboldt et al. 2010; Ross et al. 2011; Frokjaer-Jensen 2013). Indeed, this species is now an active target of research into the molecular basis of trait variation and adaptation (Baird et al. 2005; Prasad et al. 2011; Ross et al. 2011; Stegeman et al. 2013), the evolution of development (Delattre and Felix 2001; Hill et al. 2006; Guo et al. 2009; Marri and Gupta 2009), and speciation (Woodruff et al. 2010; Baird and Stonesifer 2012; Kozlowska et al. 2012; Yan et al. 2012). And yet, the limited genomic scope of understanding for its natural variation constrains our ability to fully exploit it. Here we provide the population genomic framework for relating evolutionary pressures and demographic histories to their genomic signatures in a global sample of *C. briggsae*.

## Results

### Low recombination regions show drastic skews in polymorphism

We sequenced 37 wild isolate genomes of *C. briggsae* to high coverage (median 32×; Supplementary Table 1) and identified a total of 2.70 million single nucleotide polymorphisms (SNPs) and 329 thousand short indel variants. Because of the high rate of self-fertilization in this species, each strain’s genome sequence is effectively homozygous and provides a single haploid genome. We first focused our analyses on the sample of 26 strains (including the reference strain genome) from the so-called ‘Tropical’ circumglobal phylogeographic group (Supplementary Table 1), which excludes some genetically distinct strains also derived from low-latitude locations (Cutter et al. 2006; Felix et al. 2013). Polymorphisms in this Tropical sample showed a >4-fold enrichment on the chromosome arms compared to the central and tip chromosomal domains (Figure 1). Both arm and center regions are euchromatic, but arms experience higher meiotic recombination rates and lower densities of genes (Supplementary Figure 1) (Stein et al. 2003; Cutter and Choi 2010; Ross et al. 2011). Moreover, SNPs in the central chromosome domains show a greater skew toward rare variants in the population than do SNPs in the high-recombination rate arm regions (Figure 1). We quantified this skew in variant frequencies with Tajima’s D (Figure 1), the values of which fluctuate around the neutral expectation throughout much of the chromosome arm regions whereas the chromosome centers approach the minimum possible value given the sample size (Supplementary Figure 2). These patterns of polymorphism implicate selection having eliminated linked polymorphism across megabase spans and disproportionately so in low-recombination portions of chromosomes (Sella et al. 2009; Cutter and Payseur 2013).

**Figure 1.**
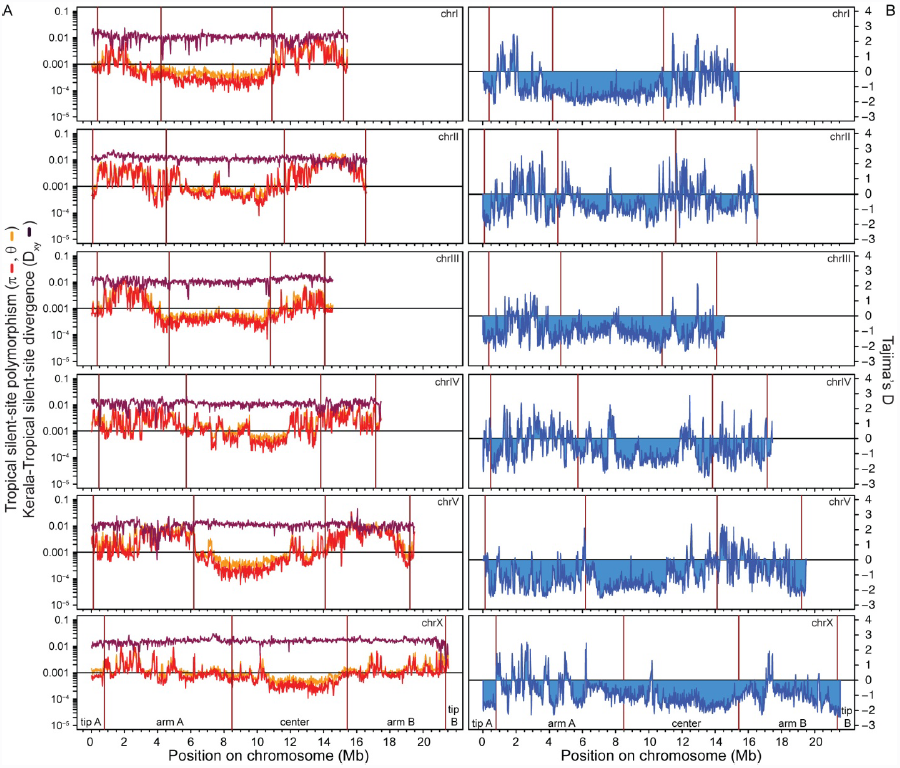
Patterns of genetic diversity depend on genomic architecture. (A) Nucleotide polymorphism of silent-sites in 20kb windows is depressed in chromosome centers (and tips) compared to arm domains for Tropical strains (median of 20kb windows on arms π_sil_ = 0.165%, centers π_sil_ = 0.041%, Wilcoxon χ^2^ = 1996.3, P<0.0001). However, absolute divergence (D_xy_) between Tropical and the distant Kerala group strains is largely insensitive to chromosomal domain. The recombination-associated domain structure of the X-chromosome is less pronounced than for autosomes (Cutter and Choi 2010; Ross et al. 2011). (B) Chromosome centers also show more skew in the site frequency spectrum, indicative of an excess of rare alleles, which is expected to result from the interaction of selection and linkage. These qualitative patterns are consistent with analysis restricted to synonymous sites (Supplementary Figure 10), indicating that differential constraint in non-coding regions does not drive the observed genomic patterns of polymorphism.

To further understand these molecular evolutionary patterns, we quantified the genomic distribution of divergence of the Tropical population relative to the ‘Kerala’ strains that are thought to have a relatively ancient split within *C. briggsae* (Cutter et al. 2010). We estimate with a molecular clock the time to the most recent common ancestor of known *C. briggsae* strains to be approximately 2 million generations ago (∼200 thousand years, assuming 10 generations per year). Using Kerala strains to infer ancestral and derived alleles in the Tropical sample allowed us to screen chromosomes for regions with an unusual incidence of new derived variants, which is indicative of recent positive selection (Fay and Wu 2000; Zeng et al. 2006). Interestingly, despite the overall skew toward rare variants in chromosome centers (Figure 1), centers did not show a biased incidence of high-frequency derived variants (Supplementary Figure 2). This result suggests that the genomic distributions of polymorphism in *C. briggsae* might more likely derive from linked selection associated with removal of deleterious mutations (‘background selection’ (Charlesworth et al. 1993; Fay and Wu 2000)).

In contrast to patterns of polymorphism, we found that absolute sequence divergence between Tropical and Kerala strains does not differ markedly among chromosomal domains (Figure 1). We can ask, however, whether the slight elevation in divergence on the arms could be explained by incomplete lineage sorting of ancestral polymorphism (median of 20kb windows of silent sites on arms D_xy_ = 0.0122, centers D_xy_ = 0.0113). Presuming that the highly selfing *C. briggsae* common ancestor had equivalent levels of polymorphism as extant Tropical strains (median of 20kb windows on arms π_si_ = 0.00165, centers π_si_ = 0.000409), divergence in chromosome centers would actually be predicted to be 10.2% less than on arms relative to the observed 7.6% reduction, suggesting that *C. briggsae* populations might have had moderately greater diversity in the past (see below PSMC analysis for evidence supportive of this hypothesis) (Supplementary Figure 3). These findings are consistent with the lack of evidence for differences in mutational input for chromosome arms and centers, based on mutation accumulation experiments (Denver et al. 2009; Denver et al. 2012), and is inconsistent with recombination-associated mutation (Lercher and Hurst 2002; Cutter and Payseur 2013). Interestingly, some chromosomes’ arms show lower divergence than centers using relative metrics (i.e. F_ST_; Supplementary Figure 4), in contrast to the opposite pattern with absolute divergence metrics, implicating differential gene flow and/or incomplete lineage sorting across the genome (Pease and Hahn 2013; Cruickshank and Hahn 2014). Altogether, we conclude that patterns of polymorphism and divergence for *C. briggsae* are most consistent with selection at linked sites having induced the distinct molecular evolutionary signatures in low-recombination centers relative to high-recombination arms.

Within the Tropical group strains, the chromosomal extent of linkage disequilibrium (LD) in *C. briggsae* is less extreme than in 39 strains of *C. elegans* (Figure 2). Nevertheless, LD in *C. briggsae* occurs between polymorphisms on different chromosomes to a greater extent than expected from sample size alone (Figure 2), implicating extreme self-fertilization as the dominant mode of reproduction in nature and consistent with a genetically effective outcrossing rate <0.0011 per generation provided N_e_ > 10,000 (Supplementary Figure 5).

**Figure 2.**
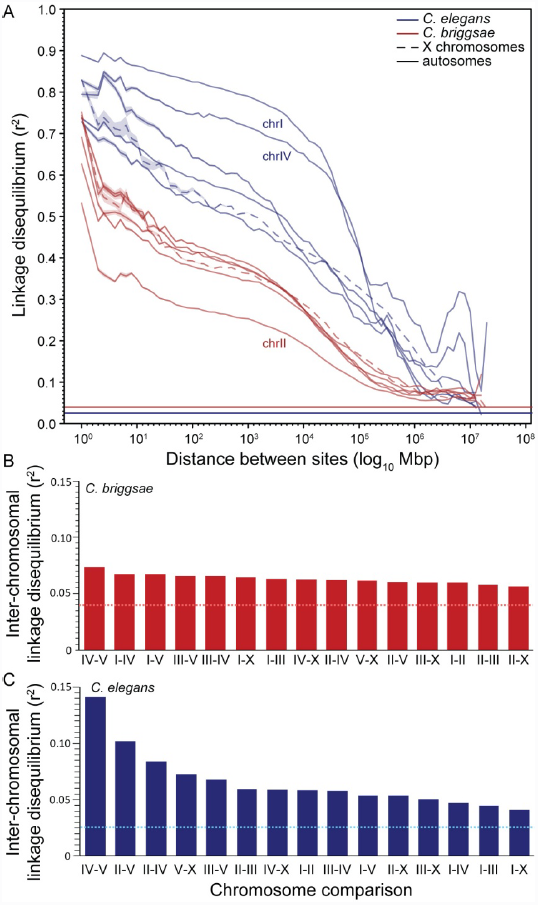
The decay of linkage disequilibrium (r^2^) is more rapid in *C. briggsae* than *C. elegans* along every chromosome. The sex chromosome is not distinctive relative to autosomes in terms of linkage disequilibrium (LD) decay in either species (A), although *C. elegans* chromosomes I and IV have elevated LD and *C. briggsae* chromosome II shows reduced LD compared to other chromosomes. The inter-chromosomal LD for *C. briggsae* (B) spans a narrower range of mean values among chromosome pairs than *C. elegans* (C), although both species have more LD between chromosomes than expected (horizontal lines). Horizontal lines indicate the background LD expected given the sample size (Weir and Hill 1980). *C. briggsae* strains include 25 Tropical strains (excluding reference strain AF16); *C. elegans* includes 39 strains (excluding Hawaiian CB4846). LD calculations exclude singleton polymorphisms.

### A diaspora of C. briggsae populations

We next constructed neighbor networks to visualize genetic distances using a set of 439,139 SNPs with complete information across all 38 strains (Figure 3). We assessed population differentiation more formally with ADMIXTURE (Alexander et al. 2009) and identified up to 8 possible subpopulations corresponding to genetic clusters observed in the neighbor network, with K=4 minimizing the cross-validation error (Figure 3). Moving from K=2 to 8, first the large sample of Tropical strains are distinguished from all others, after which the Kerala samples are revealed as distinct, followed by the Taiwanese strain pair, with further subdivisions until K=8 that separate the Montreal, Hubei, Nairobi strains, as well as splitting the Tropical strains into two subgroups (Supplementary Figure 6). The strains from Hubei-Montreal-Nairobi-Taiwan do not show evidence of simply being a group derived from admixture between the Temperate and Tropical clade strains, based on non-significance for the f_3_ test of (Reich et al. 2009) (f_3_=0.009897 ±1SE=0.000741, Z=13.35). Notably, chromosomes differ in the number of populations that minimize the cross-validation error identified by ADMIXTURE (K=3 for chromosomes 1, 3, 4, 5; K=4 for chromosomes 2, X and the full genome) (Supplementary Figure 6), suggesting varying degrees of gene flow for different chromosomes. A caveat to the ADMIXTURE analysis is that the strong linkage disequilibrium across the *C. briggsae* genome will not be properly accounted, and several potential genetic populations are represented by only one or two strains; this may explain the grouping together of the Nairobi, Montreal, Hubei, and Taiwan strains that are separated by long genetic distances (Figure 3). The pattern of genomic haplotype chunk sharing between strains, as identified with Chromopainter (Lawson et al. 2012; Yahara et al. 2013), recapitulates these trends of genetic differentiation (Figure 3). Further analysis using TreeMix (Pickrell and Pritchard 2012) implicates plausible cases of gene flow between the genomes of geographically and genetically distinct strains, although the topological position of highest-weight migration events suggest the possibility that this method detects incomplete sorting of ancestral polymorphism (Figure 3; Supplementary Figure 7). These diverse genome-wide analyses all are consistent and extend the findings from phylogeographic studies of small numbers of loci (Cutter 2006; Cutter et al. 2010; Raboin et al. 2010; Felix et al. 2013), reinforcing the identity of the Temperate and Tropical phylogeographic groups, clarifying the genomic make-up of geographically restricted strains, and underlining the great genetic distance of basal strains from Kerala, India to other *C. briggsae*.

**Figure 3.**
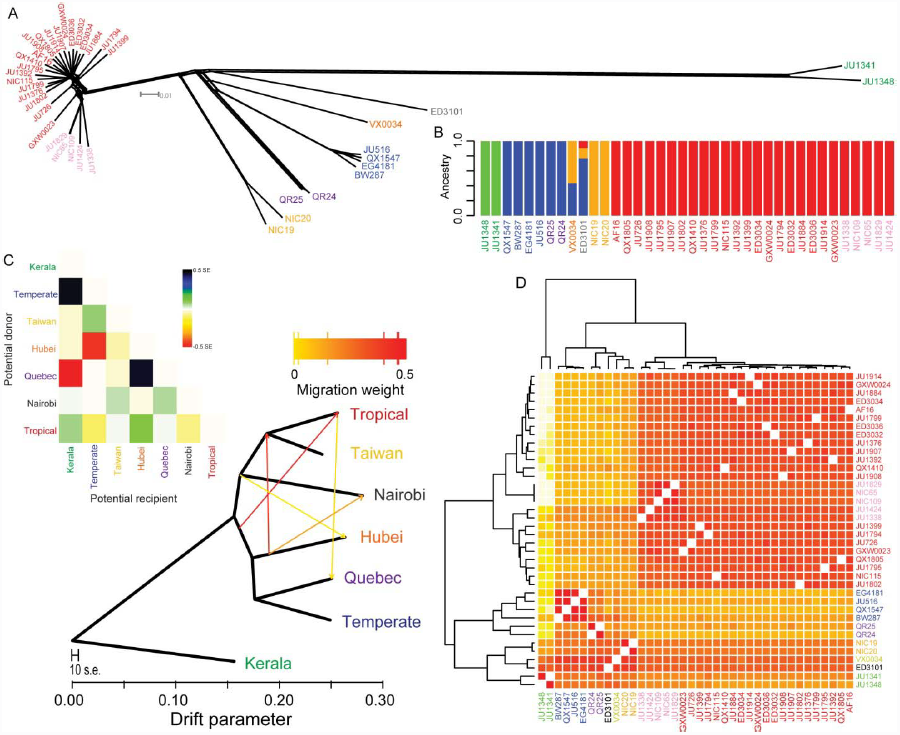
Diverse genomic analyses affirm the genetic distinctiveness of phylogeographic groups within *C. briggsae*. A neighbor network for all chromosomes (A) discriminates phylogeographic groups of strains, corresponding to the pan-global ‘Tropical’ clade (red and pink strain labels), pan-global ‘Temperate’ clade (blue), and genomic haplotype groups that exhibit restricted geographic origins around the globe (‘Quebec’ purple, ‘Nairobi’ black, ‘Hubei’ orange, ‘Taiwan’ yellow, ‘Kerala’ green). The Admixture program minimizes the cross-validation error of ancestral relationships when it identifies four genetic clusters in this dataset (B). Permitting migration in the genomic ancestry of the 37 *C. briggsae* strains with TreeMix suggests multiple plausible instances of migration (C), although incomplete sorting of ancestral polymorphism provides an alternate interpretation. Heatmap above the genealogy indicates residual fit to a model with 5 migration events. (D) Haplotype clustering of the phylogeographic groups is recapitulated in a similar manner in Chromopainter’s genome-wide co-ancestry matrix (Euclidean log2). Dendrogram on left indicates strain clustering with the unlinked model; top dendrogram indicates strain clustering with the linked model. All analyses used the set of 439,139 SNPs with allele information present in all strains.

Our sample of 4 strains from the ‘Temperate’ circum-global phylogeographic group of *C. briggsae* showed 2-to 3-fold lower polymorphism across the genome than the Tropical population sample, consistent with previous findings based on a few loci (Cutter et al. 2006). As observed for the Tropical strains, diversity for the Temperate group is reduced in chromosome centers compared to arms (median of 20kb windows on arms π_si_ = 0.00053, centers π_si_ = 0.00027; Supplementary Figure 4). We next quantified genetic differentiation between the Temperate and Tropical samples along their chromosomes. In contrast to our observations for absolute divergence, relative measures of differentiation (F_ST_) indicate stronger differentiation in chromosome centers (Figure 4, Supplementary Figure 4). These findings are fully consistent with the effects of linked selection on genomic regions with high gene density and low recombination rates (Charlesworth et al. 1997; Charlesworth 1998) and corroborate observations outside of nematodes, including humans and other mammals, birds, insects, and plants (Keinan and Reich 2010; Geraldes et al. 2011; Nachman and Payseur 2012; Pease and Hahn 2013; Renaut et al. 2013; Cruickshank and Hahn 2014), as well as the prediction that the genomes of selfing species will be particularly affected by linked selection (Charlesworth et al. 1997).

**Figure 4.**
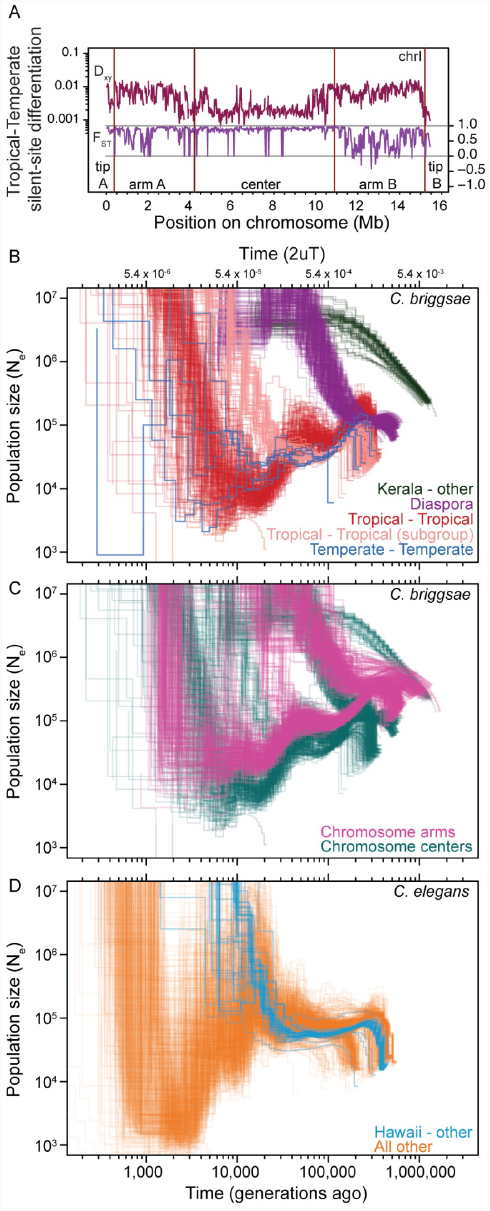
Demographic history of *C. briggsae* and *C. elegans* populations. (A) Relative measures of population differentiation (F_ST_) are greater in chromosome centers between Temperate and Tropical phylogeographic groups. By contrast, the D_xy_ measure of absolute divergence between populations shows the opposite trend, indicative of selection at linked sites being stronger in the low recombination chromosome centers (Pease and Hahn 2013; Cruickshank and Hahn 2014). Windows of 20kb for silent-sites along chromosome I is shown as an exemplar of all chromosomes (Supplementary Figure 2). Iterated pairwise sequential Markovian coalescent (PSMC) analysis of all *C. briggsae* (B-C) and *C. elegans* (D) genomes show the history of population size change and population splitting. Each line represents the change in population size (N_e_) through time inferred for a pair of genomes, with all pairs of haploid genomes among the 37 strains of *C. briggsae* and 40 strains of *C. elegans* superimposed to indicate biological replication in the inference of demographic patterns. PSMC curve profiles restricted to the upper right in (B) illustrate the deep divergence of Kerala strains to all others (∼2 million generations ago, green) and the more recent diaspora of several genetically distinct strain groups from each other nearly simultaneously 300-500 Kgen ago (purple). PSMC profiles within each of the Tropical (red, pink) and Temperate phylogeographic groups show N_e_ fluctuations in their past, with larger N_e_ in the distant past and a recent population split within the Tropical group (pink vs. red). Only analyses of chromosome center domains are shown in (B). PSMC profiles of *C. briggsae* strain pairs from within a phylogeographic group other than Temperate and Tropical are not shown. Rapid recent time N_e_ increases likely reflect an artifact of the PSMC algorithm in estimating N_e_ on short timescales (Li and Durbin 2011). (C) PSMC profiles involving Tropical strain comparisons with all other strains partitioned according to chromosomal domain (cyan center domain, magenta arm domain). Low recombination chromosome centers have lower N_e_ and more recent coalescence, and the ancestral polymorphism that generates heterogeneity in the PSMC profiles of the ‘diaspora’ differentiation of phylogeographic groups 300-500 Kgen ago is more constricted for chromosome centers. (D) PSMC analysis of *C. elegans* indicates a split of the Hawaiian CB4856 strain (blue, 30-50 Kgen ago) with all other strains in the sample (orange), and an overall strong decline in population time since then. Analyses of chromosome centers are shown, with analysis of arm regions in Supplementary Figure 8. Excludes 14 of 703 *C. briggsae* strain pairs and 10 of 780 *C. elegans* strain pairs owing to spurious PSMC profiles.

### Genomic histories of population size and differentiation are sensitive to selection and linkage

To investigate the population history of the Temperate and Tropical populations in relation to representatives from geographically restricted genetic groups, we applied the pairwise sequential Markovian coalescent (PSMC) model of (Li and Durbin 2011). We iteratively created 703 ‘pseudo-diploid’ genomes involving all pairwise combinations of the 38 strains (including the reference strain genome) to evaluate biological replication of the coalescent histories inferred by PSMC. Because of the strong recombination domain structure apparent in the *C. briggsae* genome, we performed analyses separately for arm and center chromosomal regions. The PSMC analysis revealed that the low-recombination centers of chromosomes have had a smaller effective population size (N_e_) throughout their history and coalesce in the more recent past than chromosome arms (Figure 4). This is consistent with our finding of greatly reduced polymorphism owing to linked selection in the central, low-recombination portions of chromosomes (Figure 1).

PSMC profiles for within-population genome pairs of Temperate and Tropical strains indicate elevated population size in the distant past (40-70 thousand generations ago), with low population sizes (N_e_ < 20,000) in the more recent past (10-30 thousand generations ago) (Figure 4; Supplementary Figure 8). The PSMC profiles ostensibly show rapid population growth within the last ∼8 thousand generations, but this likely represents an artifact of PSMC having difficulty estimating N_e_ on very recent timescales (Li and Durbin 2011). As anticipated from patterns of polymorphism, PSMC indicates that the Tropical effective population size has been larger than the Temperate population through much of their histories (>30 thousand generations ago; Figure 4).

We also used PSMC profiles to draw inferences about the timing of lineage splitting for geographically-distinct genetic groups of *C. briggsae*. Lineage splitting can be inferred from the pseudo-diploid PSMC profiles as a signal of rapid population size increase in the distant past, as the distinct genomic haplotypes from separated populations accrue mutations independently (Li and Durbin 2011). Our pseudo-diploid genome combinations of genetically-distinct strains revealed i) an ancient split of strains from Kerala, India (>2 million generations ago), ii) a nearly simultaneous split of 6 distinct lineages occurring approximately 200-300 thousand generations ago (Temperate, Tropical, and strains sampled in Montreal, Hubei, Taiwan, and Nairobi), and iii) a relatively recent separation of the Tropical strains into two subgroups 50-100 thousand generations ago (Figures 3, 4). Theory predicts that the low polymorphism chromosome centers should exhibit reduced heterogeneity in the timing of divergence among populations, owing to more complete lineage sorting (Cutter 2013; Pease and Hahn 2013). Indeed, inter-population divergence of the chromosome center versus arm PSMC profiles are consistent with this expectation (Figure 4; Supplementary Figure 8) (Cutter and Choi 2010; Cutter 2013; Pease and Hahn 2013). If *C. briggsae* populations pass through 10 generations per year on average, then much of this dynamic world-wide intraspecific history that is detectable in the genome has taken place within just the last few hundred thousand years.

For comparison to the striking signatures of population structure in *C. briggsae*, we analyzed the PSMC profiles for all pairs of pseudo-diploid genomes for 40 wild strains of *C. elegans* (Thompson et al. 2013) that exhibit little global phylogeographic structure (Sivasundar and Hey 2003; Barrière and Félix 2005; Cutter 2006; Andersen et al. 2012). This analysis revealed a pattern of a modestly increasing population size for *C. elegans* during its ancient coalescent history, then having suffered a strong decline in effective size recently over the past ∼10 thousand generations and experiencing a demographic split ∼30 thousand generations ago of the Hawaiian strain CB4856 from other wild isolates in the sample (Figure 4; Supplementary Figure 8). Like *C. briggsae*, chromosome centers in *C. elegans* show markedly depressed N_e_ and depth of coalescence compared to chromosome arms (Supplementary Figure 8), indicative of the influence of especially strong linked selection in the gene dense and low recombination centers of chromosomes (Cutter and Payseur 2003; Rockman et al. 2010; Andersen et al. 2012).

### Signatures of hyperdiversity in the dioecious ancestor

To further explore molecular evolutionary patterns in chromosomal centers compared to chromosomal arms, we quantified sequence divergence between species. No published genome annotation exists for *C. nigoni*, the sister species to *C. briggsae* (Kiontke et al. 2011; Felix et al. 2014). Therefore we computationally extracted and aligned a set of 6,435 ortholog coding sequences for *C. nigoni* from draft genome sequence (Kumar et al. 2012), from which we computed rates of sequence divergence at synonymous sites (dS) and replacement sites (dN) (Supplementary Figure 9). Synonymous-site divergence between *C. briggsae* and *C. nigoni* averages 20.7%, and we estimate that *C. briggsae* shares a most recent common ancestor with *C. nigoni* approximately 35 million generations ago (3.5 million years, assuming 10 generations per year).

We found that synonymous-site inter-species divergence differs significantly among chromosomes (F_5,5085_ = 72.0, P < 0.0001), with the X-chromosome and chromosome V having significantly higher rates of divergence than other chromosomes (Tukey’s post-hoc tests). The X-chromosome also shows significantly higher divergence than autosomes for distant populations of *C. briggsae* (Tropical-Kerala silent-site D_xy_; F_5,5256_ = 294.7, P < 0.0001, Tukey’s post-hoc tests), suggesting that the mutation rate of the X-chromosome exceeds the autosomes.

Curiously, genes in high-recombination chromosome arms exhibit higher dS than for genes in low-recombination centers (dS = 0.259 vs. 0.186; F_1,4854_ = 766.8, P < 0.0001; Figure 5), which also was observed for the deeper divergence between *C. briggsae* and *C. elegans* (Cutter and Payseur 2003). Three scenarios could explain this disparity in synonymous site divergence between recombination domains: i) stronger selection for biased codon usage that depresses dS among genes found in chromosome centers (Hershberg and Petrov 2008; Plotkin and Kudla 2011), ii) recombination-associated mutation (RAM) could elevate divergence in high-recombination regions (Lercher and Hurst 2002; Hellmann et al. 2003), or, iii) stronger linked selection in low-recombination regions could lead to less ancestral polymorphism (AP) for a hyperdiverse ancestral species (Begun et al. 2007; Cutter and Choi 2010; Cruickshank and Hahn 2014), because observed neutral divergence is the sum of new mutational differences since speciation and the time to coalescence for polymorphisms in the ancestral species (Gillespie and Langley 1979). Codon bias is significantly stronger for genes in chromosome centers, but by only a small magnitude (mean ENC_arm_ = 50.03, ENC_center_ = 49.11; P<0.0001). Current experimental data from *Caenorhabditis* does not support RAM (Denver et al. 2009; Denver et al. 2012), and divergence between distant populations within *C. briggsae* shows no similarly strong effect (Figure 1, Supplementary Figure 3). For the AP explanation to hold, we calculated that ancestral diversity would need to be reduced by >40% in chromosome centers, given ancestral hyperdiversity of 6-8% at synonymous sites on chromosome arms (Supplementary figure 9) (Cutter et al. 2013). *Drosophila* estimates suggest >75% reduction in diversity across the genome owing to linked selection (Elyashiv et al. 2014), but empirical estimates in outbreeding *Caenorhabditis* are not yet available. It also remains possible that differing mutational or gene conversion dynamics of the recombination domains for selfing and outcrossing species could contribute to these observations, or, chromosome rearrangements between species or the substantial genome reduction in *C. briggsae* (Thomas et al. 2012) could conspire to produce spurious ortholog inferences and artifactually strong disparity in dS between chromosome domains.

**Figure 5.**
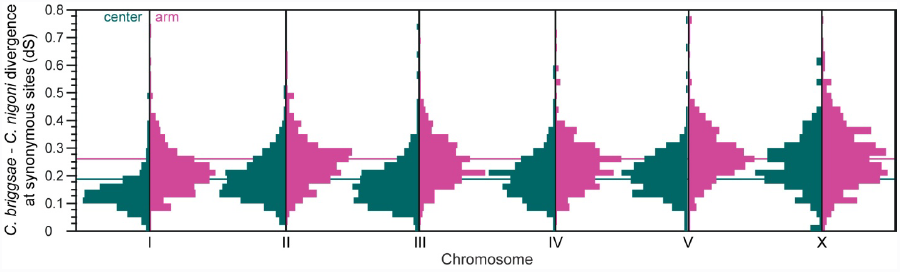
Divergence at synonymous sites for orthologs of *C. briggsae* and *C. nigoni* is higher for genes linked to arm domains than center domains, for all chromosomes (all Wilcoxon test P≤0.0035). The chromosomes also differ significantly from each other in average substitution rates (F_5,5085_ = 72.0, P < 0.0001), with chromosome I being lowest and chromosomes V and X being highest (Tukey’s HSD). Horizontal lines indicate mean dS for genes in center domains (cyan, dS = 0.186) or arms (magenta, dS = 0.259) across all chromosomes. Loci with strong codon bias (ENC < 45) excluded from analysis.

### Relaxed selection from high selfing is recorded in functional variation

In our panel of 38 *C. briggsae* genomes, we identified 209,482 SNPs that alter the amino acid sequence encoded by 18,697 genes. In addition, of the 329,281 small indel variants (1 to 60 bp long) that we identified, 10,250 of them occur in coding sequences to affect 6,879 genes. Premature stop codons often arise in these genes as a result of frame-shift or non-frame-shift indel changes (5,856 genes), and 320 genes having splice junction-spanning indels. Moreover, there are 1,244 nonsense SNP variants that create premature stop codon (PSC) alleles in 1,027 genes, in addition to those genes with premature stops induced by indel variants (1,736 genes have both indel-and SNP-induced PSCs). We also identify 356 genes with SNPs that change the stop codon in the reference genome gene annotation into an amino acid codon (stop codon losses, SCL); given that the stop codon allele is usually extremely rare or is unique to the reference sequence (Figure 6), we conclude that such sites identify premature stop mutations (or errors) in the reference gene annotation and that the true wildtype coding sequences extend farther downstream. In all, we find 18,697 of the 19,884 genes detected in the genome to harbor natural mutations that could alter the function of the encoded protein, complementing a similar resource for *C. elegans* (Thompson et al. 2013), to provide a valuable catalog of mutants for experimental analysis.

**Figure 6.**
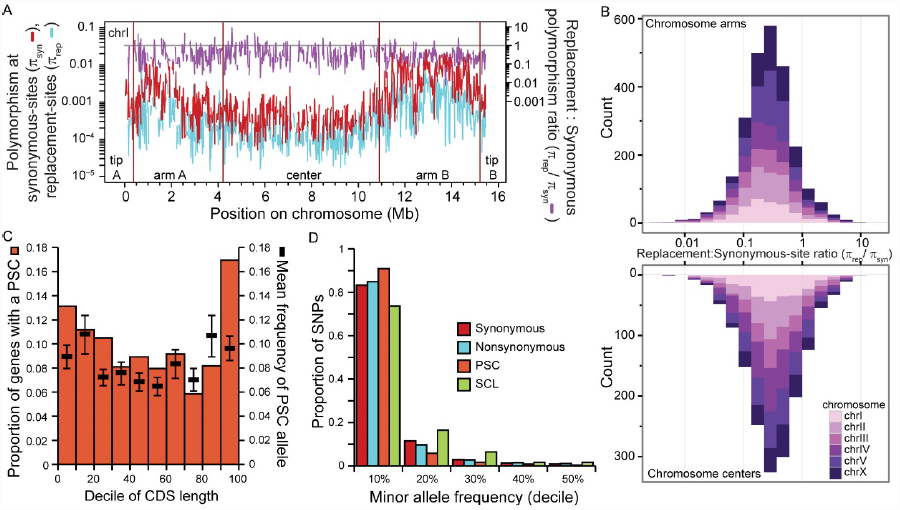
Missense and nonsense mutations are generally deleterious and selected against. (A) Windows of 20kb sliding along chromosome I indicate the lower polymorphism at non-synonymous sites (π_rep_) than at synonymous sites (π_syn_) for Tropical strains (see Supplementary Figure 10 for all chromosomes). A slight trend of elevated π_rep_/π_syn_ in chromosome centers (B) is indicative of less effective selection in purging deleterious mutations from these regions of high linkage (median of 20kb windows on arms π_rep_/π_syn_ = 0.259, centers π_rep_/π_syn_ = 0.290; Wilcoxon χ^2^=15.84, P<0.0001). The stacked histogram of π_rep_/π_syn_ in (B) shows the cumulative abundance of 20kb windows with a given bin of π_rep_/π_s_ across the 6 chromosomes, partitioned into chromosome arm and center domains. (C) Distribution of nonsense SNPs along coding sequences (CDS), expressed in percent of the CDS length. (D) Distribution of the minor allele frequency (MAF) for different classes of SNPs: synonymous (red), nonsynonymous (blue), premature stop codons (PSC, orange), stop codon losses (SCL, green). PSC SNPs have significantly skewed low MAF values, indicating stronger selective constraints. In contrast, SCL SNPs have significantly higher MAF suggesting misannotation of the reference stop codon or PSC mutations in the reference genome.

Our analysis of SNPs that create premature stop codons (PSC) indicates that such nonsense mutations are generally deleterious and disfavored by selection within wild contemporary *C. briggsae* populations. Specifically, nonsense alleles occur at significantly lower population frequency than do the minor alleles of missense or synonymous SNPs (χ^2^ = 39.94, P<0.0001; χ^2^ = 56.62, P<0.0001; Figure 6B). Moreover, PSC mutations are non-uniformly distributed along coding sequences (χ^2^ = 92.37, P < 0.0001), being more prevalent in the last 10% of coding sequences (Figure 6A), as seen in *Drosophila* (Hoehn et al. 2012; Lee and Reinhardt 2012). However, unlike in *D. melanogaster* (Lee and Reinhardt 2012), PSC-containing genes are enriched on the *C. briggsae* X chromosome (6.8% for the X vs 4.9% for autosomes, χ^2^ = 17.46, P<0.0001). Despite the general pattern of selection against PSC alleles, we find that they occur more commonly in genes that are expected to be subject to weaker selection in this species, namely, genes with low expression and genes with male-biased expression. Overall, the expression level of PSC-containing genes is 1.3-fold lower than other genes (t = 13.32, P < 0.0001) and their protein sequences evolve faster (dN/dS_PSC_ = 0.146; dN/dS_others_ = 0.106; t = 4.46, P < 0.0001). Moreover, male-biased genes contain a disproportionate load of these polymorphic nonsense mutations (χ^2^ = 6.2987, P = 0.012), indicative of relaxed selection on male functions in *C. briggsae*.

To characterize genic influences of selection across the genome, we contrasted polymorphism (π_rep_, π_syn_) within *C. briggsae* and divergence (dN, dS) relative to *C. nigoni* for nonsynonymous and synonymous coding sites (Figure 6, Supplementary Figure 10). Replacement site SNPs for the Tropical population sample are less common in chromosome centers than arms, mirroring the pattern for synonymous polymorphisms (median of 20kb windows on arms π_rep_ = 0.073%, centers π_rep_ = 0.020%, Wilcoxon χ^2^ = 1057.6, P<0.0001). This finding is reminiscent of the observation in *C. elegans* that genomic context, owing to linked selection, is a more important driver of heritable variation in gene expression than are the functional attributes of the encoded genes (Rockman et al. 2010). Indeed, sliding windows of π_rep_/π_syn_ indicate that chromosome centers have an 11% relative excess of non-synonymous SNPs, despite the paucity in absolute numbers of SNPs in centers (median of 20kb windows on arms π_rep_/π_syn_ = 0.259, centers π_rep_/π_syn_ = 0.290; Wilcoxon χ^2^ =15.84, P<0.0001; Figure 6). This relative over-abundance of non-synonymous SNPs in chromosome centers suggests that linkage interferes with the effective elimination of detrimental mutations to a greater extent in such regions. By contrast, divergence with *C. nigoni* at replacement sites does not depend on affiliation with chromosomal domains (median dN/dS arms = 0.0711, centers 0.0719; Wilcoxon *χ*^2^ = 0.172, P = 0.68), implicating recent demography and linked selection within *C. briggsae* as being responsible for differential efficacy of selection on functional protein variation among chromosomal domains. Moreover, the distribution of π_rep_/π_syn_ relative to dN/dS and π_rep_/π_syn_ in related outbreeding species indicates that selection has failed to purge many deleterious nonsynonymous-site mutations from the *C. briggsae* population (median π_rep_/π_syn_ = 0.275, dN/dS = 0.0727; mean *C. brenneri* (Dey et al. 2013) π_rep_/π_neu_ = 0.026).

## Discussion

This first full-genome characterization of *Caenorhabditis* population polymorphism reveals the striking evolutionary histories of self-fertilization, population splitting, and natural selection of *C. briggsae* and *C. elegans*. We demonstrate that natural selection and genetic linkage interact to extraordinary effect to eliminate both neutral and functional genetic variability in low-recombination regions of *C. briggsae*’s genome. Low-recombination chromosome centers are gene rich in this highly self-fertilizing species, creating the perfect storm for genetic hitchhiking and background selection, with its high density of selected targets in strong genetic linkage (Maynard Smith and Haigh 1974; Charlesworth et al. 1993; Cutter and Payseur 2013). Intriguingly, this mark of selection on the genome also reveals itself in the profiles of coalescent history for both *C. briggsae* and *C. elegans* as analyzed with the Markovian coalescent modeling framework (Li and Durbin 2011), indicating that interpretations from PSMC analyses are sensitive to effects of the interaction between selection and linkage, which may also assist in identifying histories of population splitting. Emerging techniques will be especially valuable in quantifying the relative contributions of selective sweeps for positively selected alleles and purifying selection against detrimental alleles in creating these patterns (Elyashiv et al. 2014). However, in contrast to *C. elegans* (Andersen et al. 2012), the absence of a strong signal of high-frequency derived polymorphisms in *C. briggsae* chromosome centers implicates background selection to eliminate deleterious mutations as the primary cause of genomic variation in the distribution of polymorphisms in this species (Kaiser and Charlesworth 2009; Cutter and Choi 2010; Cutter and Payseur 2013). A striking consequence of such a strong interaction between linkage and selection in the genomes of these self-fertilizing species is that evolution on recent timescales will be determined to a large extent by where in the genome new mutations happen to arise, rather than by the individual functional effects of those mutations.

Plant species also reveal dramatic influences of self-fertilization on polymorphism in their genomes (Wright et al. 2013). The less diverse genome of self-fertile *Mimulus nasutus*, compared to outbreeding *M. guttatus*, also experiences strong linkage disequilibrium and an abundance of premature stop codon alleles (Brandvain et al. 2014). The recent origins of self-fertilization in *Capsella rubella* from its progenitor *C. grandiflora*, and between *Arabidopsis thaliana* and *A. lyrata*, has yielded similar effects (Foxe et al. 2009; Cao et al. 2011; Brandvain et al. 2013). Unlike these species, however, we find that *C. briggsae*’s euchromatic genome architecture that combines high gene density and low recombination rate lend it especially pronounced signatures of linked selection, with the effect on *C. elegans*’ genome even more extreme (Andersen et al. 2012). Whether the corresponding genomic regions in obligatorily outbreeding, often hyperdiverse, species of *Caenorhabditis* will exhibit similar evolutionary signatures remains an open question (Cutter et al. 2013; Cutter and Payseur 2013): to what extent will more effective recombination from outbreeding free the genome from the effects of linked selection, and might soft sweeps play an increased role in shaping genome evolution in such species? General answers to questions like these will benefit from the merger of population genomic analysis with phylogenetic comparative methods.

The globally-distributed *C. briggsae* is comprised of a diaspora of differentiated lineages, most of which separated from one another ∼200 thousand generations ago (∼20 Kya). Our iterated pairwise sequential Markovian coalescent analyses complemented more conventional procedures for inference of *C. briggsae’s* evolutionary history to reveal the timing of differentiation and population size change. We find little evidence for ongoing gene flow among these lineages, although most of the distinct lineages have been sampled only rarely in nature, with the ‘Temperate’ and ‘Tropical’ genetic groups predominating in wild collection efforts (Felix et al. 2013). Moreover, the deep split of ‘Kerala’ strains from all other *C. briggsae* lineages ∼2 million generations ago (∼200 Kya), averaging 1.2% silent-site sequence divergence, is an especially striking feature of the evolutionary history of this species. This splitting reinforces the notion that *C. briggsae* provides a powerful developmental genetic system for studying adaptive divergence and incipient speciation, particularly when combined with evidence of differential adaptation to temperature by Temperate and Tropical genotypes (Prasad et al. 2011), indications of some post-zygotic fitness barriers (Dolgin et al. 2008; Ross et al. 2011; Baird and Stonesifer 2012), and with available genetic toolkits (Koboldt et al. 2010; Ross et al. 2011; Frokjaer-Jensen 2013) and the genomic resource provided here.

In addition to revealing these evolutionary phenomena, the >3 million polymorphisms that we discovered in wild strains of *C. briggsae*, of which 7.25% alter the coding sequences of 94% of the genes in the genome, provide a trove of mutant alleles for functional analysis. As comparative molecular genetic and developmental studies accelerate our understanding of fundamental processes like developmental genetic networks, cell morphogenesis and cell lineage (Wang and Chamberlin 2004; Zhao et al. 2008; Riche et al. 2013; Chen et al. 2014; Ellis and Lin 2014; Verster et al. 2014), such mutational inventories provide a crucial experimental resource (Thompson et al. 2013). This study of genomic polymorphism in wild populations of *C. briggsae* provides a foundation to probe the consequences for molecular function to such genomic change and in response to divergence of traits.

## Methods

### Population genomic sequencing

We shotgun sequenced genomes to at least 15× coverage for 37 *C. briggsae* strains derived from 7 previously-hypothesized global phylogeographic groups (median 32× coverage per strain), with deepest geographic sampling to obtain 25 ‘Tropical’ group strains (Cutter et al. 2006; Cutter et al. 2010; Felix et al. 2013). Details about sequencing library and platform for Illumina sequencing for each strain are given in Supplementary Table 1, with data deposited in the NCBI Sequence Read Archive (#XXXXXX).

### Mapping and SNP determination

We performed a first-pass mapping of perfect reads using the Burrows-Wheeler Aligner v.0.2.2-r126 (Li and Durbin 2009) to the WS242 (cb4) reference genome assembly for ‘Tropical’ *C. briggsae* strain AF16 (www.wormbase.org), which also was used later for annotated feature extraction. We followed this with Stampy v.1.020 for mapping the remaining divergent reads (Lunter and Goodson 2011). Picard tools v.1.96 was used for file format manipulation to integrate with the Genome Analysis ToolKit (GATK v.2.7-4) (Van der Auwera et al. 2002). For SNP calling, we applied the GATK UnifiedGenotyper with ploidy = 1, given the highly inbred nature of *C. briggsae* strains. After filtering to incorporate per strain mapping quality and depth of coverage, we derived a total of 2,700,664 sites with single nucleotide polymorphisms (SNPs) across all strains, of which 439,139 SNPs had high-quality allele calls in every strain. We applied BreakDancer (Chen et al. 2009) and Pindel (Ye et al. 2009) to identify short indel variants ≤60 bp long, requiring ≥3 supporting reads for inclusion in subsequent analyses. Repeats were identified in the cb4 reference genome using RepeatModeler and RepeatMasker. Polymorphism data has been deposited in WormBase (http://www.wormbase.org; WBVar#XXXXXX-#XXXXXX).

### Population genetic metrics

We computed standardized per-nucleotide measures of polymorphism (π, average number of pairwise differences; θ_w_ number of segregating sites) in 5,262 non-overlapping 20kb windows using a corrected site frequency spectrum (SFS) (Nielsen 2005; Wakeley 2009; Hufford et al. 2012). These metrics were calculated separately for nonsynonymous sites, synonymous sites, and all silent sites (synonymous sites plus non-repetitive non-coding sites); only windows where at least 5000bp of informative sequence passing our filters are shown in sliding window analyses plots. For polymorphism metrics of the Tropical population sample, we required ≥20 strains to have informative sequence data for a given site. The SFS of polymorphisms from the Tropical population sample was summarized for silent sites with unfolded metrics using the divergent Kerala strains as outgroup (Fay and Wu’s H (Fay and Wu 2000)) and folded metrics that do not require an outgroup (Tajima’s D (Tajima 1989) and Schaeffer’s D/Dmin (Schaeffer 2002)). Differentiation between Tropical and Temperate population samples was computed with F_ST_ (Hudson et al. 1992). We also calculated absolute divergence between *C. briggsae* population sets with D_xy_ (Nei and Li 1979). Per-gene metrics were calculated only for genes in which at least 100bp, or 25% of the sequence length for genes whose CDS is shorter than 100bp, had passed filters for informative sites. We excluded 789 annotated genes that lacked an ATG start codon for their coding sequence, lacked a termination codon, or had a mismatch in their annotated length. Correction for multiple hits was applied according to (Jukes and Cantor 1969) and (McVean et al. 2002).

To compute linkage disequilibrium (r^2^) for *C. briggsae* with vcftools (Danecek et al. 2011), we merged all 37 vcf files and extracted the 2.7 million SNP sites into a new vcf file, which was set as containing phased haplotypes for analysis. We then computed r^2^ for the 25 sequenced ‘Tropical’ group samples, excluding singleton SNPs. This procedure was also applied to 39 *C. elegans* strains (excluding Hawaiian CB4856) from (Thompson et al. 2013). Inter-chromosomal linkage disequilibrium (LD) was calculated as the average r^2^ for all pairs of sites between a given pair of chromosomes, computed for each of the 15 pairings of the 6 chromosomes. Outcrossing rate was then computed from inter-chromosomal LD using the approach of (Cutter 2006), after first subtracting the expected value of r^2^ given a sample size of 25 or 39 for *C. briggsae* or *C. elegans*, respectively (E[r^2^] ∼ 1/n) (Weir and Hill 1980).

### Ortholog identification and divergence time calculation

Sequences for *C. nigoni* (formerly known as *C. sp*. 9) (Felix et al. 2014) were extracted from the 30 November 2012 assembly from (Kumar et al. 2012) (http://wormgenomes.caltech.edu). We ran tblastn v. 2.2.26+ against the *C. nigoni* genome using the *C. briggsae* CDS transcripts (version WS242 from wormbase.org), and then extracted the best-hit regions in the *C. nigoni* genome, requiring >80% overlap and collinearity to the corresponding *C. briggsae* gene, as well as requiring >80% sequence identity to exclude contaminating sequence from *C. afra* that is known to be present in the draft *C. nigoni* assembly (E. Schwarz, pers. comm.). The aligned regions of each collinear best hit were then concatenated into a preliminary *C. nigoni* gene sequence, which we subjected to blastx using *C. briggsae* WS242 protein sequences to obtain gene pairs with mutual best hits between *C. nigoni* and *C. briggsae* for use in our 6435 ortholog gene set.

We computed an estimate of the divergence time between *C. briggsae* and *C. nigoni* according to *T* = (dS – π_anc_)/(2μ) (Gillespie and Langley 1979), where dS_arm_ = 0.259, assuming silent-site polymorphism on the arms of the dioecious common ancestor is π_anc-arm_ = 0.07 (A.D. Cutter, unpublished data), and a per-site mutation rate each generation of μ = 2.7 × 10^-9^ (Denver et al. 2009). This yields *T* = 35.0 Mgen ago; lower π_anc_ or μ would result in more ancient *T*. Divergence in low recombination center regions of chromosomes might better reflect the time since the cessation of gene flow between species, owing to reduced ancestral polymorphism (Cutter 2013; Cruickshank and Hahn 2014), so we also computed *T* given dS_center_ = 0.186 and assuming π_anc-center_ = 0 to yield *T* = 34.8 Mgen ago; higher π_anc_ or μ would result in more recent *T*. These estimates of neutral divergence excluded loci with dS > 0.8 and ENC < 45 to mitigate against any remaining false ortholog assignments and genes with strong selection on codon usage. For divergence time estimation of the Kerala strains to other strains of *C. briggsae*, we used this same approach using silent-site D_xy_ instead of dS, with values of D_xy-center_ = 0.0113 and π_anc-center_ = 0.00041 to yield *T* = 2.02 Mgen ago.

### Analysis of nonsense alleles

Analyses of premature stop codon (PSC) mutations within coding sequences (CDS) excluded genes containing indel-induced premature stops. For the corresponding 1,027 genes, when multiple nonsense mutations occurred in a gene, we determined the most 5’ PSC incidence along the CDS in decile bins of length in order to capture the degree of CDS truncation. Divergence analyses between *C. briggsae* and *C. nigoni* excluded 29 genes with dS = 0 and 24 genes with dN/dS > 1. To estimate overall gene expression, we used the average expression level measured across 10 embryonic time points in strain AF16 from (Levin et al. 2012). Sex-biased expression was inferred from expression levels of AF16 males and *she-1* pseudofemales in (Thomas et al. 2012).

### Phylogeographic and demographic analysis

We performed phylogeographic analyses using the subset of 439,139 SNPs with perfect information in all strains. Splitstree (Huson and Bryant 2006) was run separately for each chromosome as well as for a concatenation of sites from all chromosomes to create neighbor-network trees for strain genomes, with reticulation in the network indicating recombination and gene flow. We next ran Admixture on the samples (Alexander et al. 2009), varying population number K from 1 to 9 for data from each chromosome separately or all together. We applied the Chromopainter program (Lawson et al. 2012; Yahara et al. 2013) to cluster strains. For this analysis, we set the recombination scaling start value to 0.2 and input the recombination rate of polymorphic sites based on linear interpolation of genetic and physical distances as reported previously (Cutter and Choi 2010; Ross et al. 2011). Based on the results of these previous analyses, for input into Treemix (Pickrell and Pritchard 2012), we grouped the 38 strains (37 strains sequenced here plus reference genome of AF16) into 7 groups (Supplementary Table 1), using the two Kerala strains as outgroup. We ran Treemix allowing 1 to 5 migration events, using groups of 100 SNPs to account for linkage disequilibrium.

We conducted historical demographic analysis with PSMC (Li and Durbin 2011), considering each strain as a single genomic haplotype. We constructed pseudo-diploid genomes for all 703 possible combinations of the 38 strains and restricted SNP identity to the set of 2.7 million SNPs. We excluded 14 pseudo-diploid strain combinations from further analysis owing to spurious PSMC profiles. PSMC was run separately for arm and center chromosomal domains as defined in (Ross et al. 2011). We ran the same pipeline on 40 *C. elegans* strain whole genome sequences and sub-chromosomal domains (Thompson et al. 2013). Out of the 780 pseudo-diploid *C. elegans* strain combinations, 10 were excluded owing to spurious PSMC profiles.

## Acknowledgements

This research was supported by funds to A.D.C. from the Natural Sciences and Engineering Research Council of Canada, a Canada Research Chair, and the National Institutes of Health (GM096008).

